# BioDCASE: Using data challenges to make community advances in computational bioacoustics

**DOI:** 10.64898/2026.04.02.716062

**Authors:** Dan Stowell, Ester Vidaña-Vila, Ines Nolasco, Ben McEwen, Lucie Jean-Labadye, Yasmine Benhamadi, Gabriel Dubus, Benjamin Hoffman, Pavel Linhart, Ilaria Morandi, Dorian Cazau, Brian Miller, Elena Schall, Clea Parcerisas, Anatole Gros-Martial, Ilyass Moummad, Pierre-Yves Raumer, Ellen White, Paul White, Paul Nguyen Hong Duc, Vincent Lostanlen

## Abstract

Computational bioacoustics has seen significant advances in recent decades. However, the rate of insights from automated analysis of bioacoustic audio lags behind our rate of collecting the data – due to key capacity constraints in data annotation and bioacoustic algorithm development. Gaps in analysis methodology persist: not because they are intractable, but because of resource limitations in the bioacoustics community. To bridge these gaps, we advocate the open science method of data challenges, structured as public contests. We conducted a bioacoustics data challenge named BioDCASE, within the format of an existing event (DCASE). In this work we report on the procedures needed to select and then conduct useful bioacoustics data challenges. We consider aspects of task design such as dataset curation, annotation, and evaluation metrics. We report the three tasks included in BioDCASE 2025 and the resulting progress made. Based on this we make recommendations for open community initiatives in computational bioacoustics.

## 1. Introduction

Computational bioacoustics has contributed much to bioacoustic study in recent years, often through the use of data-driven methods such as machine learning (Stowell, 2022). Such methods have helped to enable passive acoustic monitoring (PAM) at large scale by automated labelling of acoustic data. There has been remarkable progress, but progress remains patchy and unbalanced. Birdsong species classification has reached an impressively high level of performance across a large number of species, and is now used as a tool in many projects (Funosas et al., 2025). Yet achieving similar performance across the full range of soniferous taxa is difficult—and even with birdsong, species recognition is most reliable in Europe and North America due to the preponderance of labelled data from those regions (Funosas et al., 2025). There are also many computational tasks of interest beyond species recognition.

Innovations can originate in many different ways, but the example of birdsong species classification shows the importance of public, open community data challenges. Work on this domain accelerated strongly from around 2013, stimulated by data challenges based on open data (Fodor, 2013; Glotin et al., 2013; Goëau, Glotin, Vellinga, Planqué, & Joly, 2016; Goëau, Glotin, Vellinga, & Rauber, 2014)). Can we provide other such stimulus, to ensure that bioacoustic work of all kinds gains the maximal benefit from computational methods?

To facilitate algorithm development and evaluation, various public datasets, benchmarks and software have been published in the past five years, covering larger scales than previously (Chasmai, Shepard, Maji, & Horn, 2024; Faiß, Ghani, & Stowell, 2025; Gómez-Gómez, Vidana-Vila, & Sevillano, 2023; Pérez-Granados & Sébastián-González, 2024; van Merriënboer, Hamer, Dumoulin, Triantafillou, & Denton, 2024). There have also been many community initiatives addressing acoustic data and its analysis, such as DCLDE (Frazao, Padovese, & Kirsebom, 2020), Life-CLEF/BirdCLEF (Goëau et al., 2016, 2014), DCASE (Stowell, Giannoulis, Benetos, Lagrange, & Plumbley, 2015).

Taken together, these provide a large improvement in the resources available to anyone wishing to apply computational bioacoustics in their own project. Yet despite these advances, practitioners often encounter barriers in applying automated bioacoustics in their projects due to technical gaps that remain stubbornly un-bridged. To give some examples:

- Robust generalisation: this is a generic requirement that comes into play for almost all automatic recognition systems, especially those that are trained from data. Although there have been many advances in the size and coverage of public bioacoustic datasets in recent years, the diversity and complexity of bioacoustic soundscapes frequently ranges beyond what is captured in training data. Further, *domain shift*, the acoustic changes due to varying conditions over years/seasons/sites, implies that evaluations must explore generalisation with more than just a simple ‘test set’ held out from the training data (Nolasco et al., 2023; van Merriënboer et al., 2024).
- Spatial sound, in various aspects: from distance estimation or direction estimation, to species recognition that has clearly calibrated reliability with regard to distance. A key reason that spatial sound topics are under-addressed is that, although many large bioacoustic datasets exist, it is rare for them to include explicit spatial information such as the distance from the microphone to the animal. Another is the diversity of hardware setups that might be used e.g. for multi-microphone triangulation.
- “Tiny” systems that can run with low power demands, on small embedded devices, and/or with low ecological footprints (Millar, Haddadi, & Madhavapeddy, 2025). A recurring issue for evaluation in this domain is how to balance multiple desiderata, e.g. high recognition performance and low power consumption.
- Terrestrial and marine bioacoustics communities (as well as others, e.g. freshwater) use many of the same techniques, but often with dramatically different acoustic deployment conditions and technical requirements. The unification of computational methods across these sibling communities is desirable, especially as deep learning makes methods increasingly domain-agnostic and multi-capable. Such unification requires technical development as well as community coordination.

Each of the above tasks has received at least some attention in academic and applied research. To push forward on innovations, the generality of solutions, and the technical readiness level, experience has shown that clear comparative evaluation campaigns are valuable. To that end, in 2025 we introduced a new community evaluation campaign called BioDCASE, with its format derived from the pre-established DCASE initiative. We engaged with bioacoustic communities to determine which needs could be addressed and how, and conducted the BioDCASE 2025 evaluation initiative, to help bridge some of the gaps identified.

With this paper we make the following contributions:

1. We describe the methodology of the *data challenge*, adapted from other machine learning community initiatives and customised for bioacoustics. In particular, we consider the curation of datasets and the choice of evaluation measures, as key factors in designing tasks to represent generic but realistic bioacoustic processing needs.
2. We consider community aspects explicitly: the facilitation of bioacoustics and software communities to collaborate in this manner.
3. We report the design and results of three tasks conducted in the 2025 BioDCASE challenge: multi-channel audio alignment, detection of Antarctic blue and fin whale calls, and bioacoustics for tiny hardware.

In the following, we begin with general considerations for data challenge design. We then address the specific tasks conducted in 2025, the datasets curated for them, and the results of these evaluations. We then draw wider conclusions and recommendations for data challenges in bioacoustics. Our hope is that this paper enables the bioacoustics community to overcome common obstacles in data processing, through the communal and well-targeted use of public data challenges.

## 2. Materials and Methods

The BioDCASE challenge is a community-driven initiative where researchers from various backgrounds come together to propose and manage different bioacoustic related *tasks*. A task, in this context, is a formalised specification of a question in computational bioacoustics, which lacks a standard solution or established procedure. Tasks are thus conceptualised as an example scenario or sandbox where researchers can employ their skills and compete to create the best computational solutions to the problem, thereby contributing to accelerate innovation and development around these questions.

### Design Principles

Task design is grounded in three main principles: *Accessibility*, *Measurability* and *Generalisation*. The Accessibility principle defines that all materials are available to the wider public, where datasets, evaluation processes, and baselines are open sourced and easily accessible for download and manipulation. The Measurability principle requires that some criterion of success is available, usually a numerical performance score. This ensures that community efforts can be robustly and independently compared. The Generalisation principle is also fundamental: the choice to solve problems via public data challenges is partly motivated by the idea that the challenge can represent general issues via the specific tasks it proposes. In that sense, tasks should be considered as specific examples of broader questions, that promote the development of general solutions that can apply beyond the specific domain of the task itself. Furthermore, tasks design should also aim to represent the generality of what might be encountered in the future.

While tasks are not trivial, in the sense that there are not obvious solutions already established, they are designed to support participation of the wider research communities across the computational and bioacoustics fields, in particular entry-level researchers. Also here, the principle of accessibility plays an important role to promote participation. Before elaborating further on the details of task design, we must first set the context of the overall challenge setup.

### 2.1. Challenge and task design

Similar to the DCASE challenge (Mesaros, Serizel, Heittola, Virtanen, & Plumbley, 2025), each edition is composed by a selected set of tasks which are run independently, with unique objectives and specific datasets. All tasks develop under the same timeline, and competition outputs are gathered in the BioDCASE website. BioDCASE, as the umbrella platform for all tasks, also defines a set of general rules. These describe aspects such as how many submissions per team are allowed, requirements for the technical reports, or how evaluation data and annotations should be used. Other aspects are regulated individually by each task organising team such as the use of external data.

The challenge develops over 4 main stages, culminating on the final BioDCASE workshop day.

**Stage 0:** Challenge coordination and preparation including: request for proposed tasks, a meeting to discuss and guide shortlist of proposed tasks, selection of tasks, and preparation of datasets, rules, and guiding documents for each task. **Stage 1:** The challenge starts with the release of the development dataset, evaluation framework and baseline code. Participants start developing their own solutions.

**Stage 2:** The evaluation dataset is released, and participants submit their system outputs (e.g. predictions) calculated for the evaluation data, along with a technical report describing their approach and methodology. While source code of submitted systems is not mandatory to be made available, this is as aspect is highly valued in the subjective evaluation of the systems.

**Stage 3:** In this final stage, task organisers rank the submitted systems by running evaluation scripts that compare the produced outputs against ground-truth annotations of the evaluation set. the rankings for each task are then announced on the BioDCASE website, together with the technical reports.

**BioDCASE workshop:** Also following the structure introduced by DCASE, all tasks organising teams and participants come together in a workshop event.

**Typical Task Categories:** Tasks, in the Machine listening field, typically fit certain categories such as classification, sound event detection, generation, and clustering, among others. These categories define the core objectives of the task as well as the expected formats of system inputs and outputs. For example, classification tasks typically involve short audio segments (e.g. .wav files) paired with corresponding labels (e.g. provided in .csv or .txt files), with the goal of correctly assigning labels to previously unseen audio examples. Defining a challenge task within one of these established computational task categories is helpful for clarifying its objectives and expected outcomes. However, tasks are not limited to the standard goals of these categories. Tasks can be made more challenging by introducing additional constraints or by designing evaluation setups that emphasise specific challenging scenarios or focus on evaluation of specific capabilities of the systems. Such considerations promote the development of solutions that are applicable across a wider range of scenarios. For example, model size or computational cost may be explicitly measured and incorporated into the evaluation framework. Another common approach is the use of an out-of-domain evaluation setting, in which systems are ranked based on their performance on evaluation data drawn from a different source than the training data.

**Task Structure:** A task consists of 3 main components, each requiring careful consideration:

**Dataset:** The construction of the dataset is closely linked to the conceptualisation of the task and the desired level of difficulty. For example, an otherwise simple sound event detection (SED) task can be made substantially more challenging by incorporating noisy or acoustically complex recordings. As such, data curation is a critical step that highly influences the overall success of the task. This process involves not only assessing data quality and carefully selecting audio recordings, but also defining and applying a consistent and well defined annotation procedure. This process is necessarily made in close collaboration with the data owners and domain experts, and the tools developed to the effect or standards defined can be an important asset for the research community.

To fit the challenge setup and enable comparison across participating systems, datasets are split into fixed subsets. A development set is provided to support model training and validation, while a hidden evaluation set is used to generate final scores and rankings from trained models. In machine learning, generalisation is classically considered via the importance of maintaining independent datasets for these purposes. In that domain, the term ‘data leakage’ is used to consider issues of non-independence, such as when a random shuffle of data might split a single bout of vocalisation into one part for the training set and one part for the testing set. Thus it is important to “avoid placing different calls from the same animal encounter in both the training and test sets” (Hildebrand, Frasier, Helble, & Roch, 2022). It is further recognised that even one good test set, for final evaluation, may not be the end of the story. It is now common to evaluate algorithms on more than one test set, gathered in varying deployment conditions, to explore across how many domains the algorithm can maintain useful performance. This has been explored in prior challenges (Nolasco et al., 2023) and benchmarks (van Merriënboer et al., 2024), and will be considered in our task design.

We want to highlight that the datasets created are a valuable output of BioDCASE. They contribute directly to the definition of benchmarks, and form crucial materials supporting research outside of the challenge. In practical terms, data can be made available in platforms such as *Zenodo* ^1^ and it should be organised in ways that facilitate its download or quick inspection of its contents and format. The reference labels of the hidden evaluation set, may be made available after the challenge finishes, however that is not mandatory particularly if the task is expected to continue on subsequent editions.

Ground-truth annotations will always be needed for a task: not least for the evaluation data, and often also for training data. If these are not already available, then they must be made. The activity of preparing a challenge task often leads to contributions in data annotation and the software tools required to produce those annotations.

**Evaluation framework:** This defines how submitted solutions are assessed, compared, and ranked. In practice, the combination of evaluation metrics and the evaluation dataset determines which aspects of system performance are emphasised, which are weighted more heavily, and which may be considered less relevant or ignored.

There is no unified evaluation process shared across all tasks in the challenge. Instead, each task defines its own set of evaluation metrics that fit with the specific objectives and type of task.

During the first stage of the challenge, when participants are developing their systems, the evaluation base code is an important tool for self-assessing and guiding further development. At the end of the challenge, task organisers run the evaluation scripts on the outputs of submitted systems in order to compute the final ranking.

For each task, an additional jury award is attributed to systems that, not being necessarily high ranking, propose a new approach or are submissions of great quality and completeness.

**Baseline systems:** These provide example solutions to the problem and serve as reference points for participants. During development, participants have access to baseline performance on the validation set, information which they can use to compare against their own systems and assess progress.

Baseline systems are designed to establish a starting point and expected minimum performance level. These systems should reflect the common practices in the field, and in doing so, provide context for interpreting performance improvement and potential impact in the wider community. Indeed baseline systems are not necessarily machine learning base solutions, particularly in cases where established classical approaches exist.

An important consideration when developing a baseline is that it can work as an easy entry point for participants, therefore providing a clear and well-documented source code that participants can reproduce and improve over.

Finally, the choice of a baseline approach should be made carefully, as it may implicitly influence commonly accepted practices for similar tasks, both within and beyond the challenge context.

### 2.2. Challenge coordination

Next, we describe the coordination and organisation of the 2025 edition of the BioDCASE challenge, which was planned and coordinated by the organising committee composed of four members. Each member was assigned a role: a general chair, a workshop chair—in charge of organising the workshop—, a tasks chair—the contact point of the task organisers—, and a web chair—in charge of updating the website of both the workshop and the challenge. The four members of the committee held periodic coordination meetings since the conception of the BioDCASE initiative until the end of the edition during the workshop.

The first task of the organising committee was to select the tasks to be held. For that purpose, a public meeting was announced and promoted in October 2024 among the bioacoustics community, particularly targeting well-established research groups and senior researchers. During the meeting, the concept and objectives of BioDCASE were presented, and a brainstorming session was conducted to identify research topics of high relevance and interest to the community. At the end of the session, six topics related to bioacoustics were listed, and interested researchers willing to organise tasks related to those topics would write their names next to the ideas. As a follow-up to this meeting, a call for interest was circulated via email, in order to evaluate feasibility and data availability. After further assessment, three of the original proposals were selected and developed into official BioDCASE 2025 tasks: Task 1: Multi-Channel Alignment, Task 2: Supervised detection of strongly-labelled Antarctic blue and fin whale calls, and Task 3: Bioacoustics for Tiny Hardware. Remaining proposals were not pursued due to practical constraints such as absence of data or dedicated teams to lead their implementation.

After that, the organising committee maintained continuous interactions with the task teams, specially through the task chair, which enabled a successful delivery of the tasks as planned and announced on the website by the web chair.

The schedule of the challenge was the following: On 14 February 2025, the tasks description was announced in the official website; on 1 April 2025, the challenge launched with the release of the development datasets and baselines; on 1 June 2025, the evaluation sets were released so participants could calculate their metrics and test their models; on 15 June 2025, the challenge closed, meaning that participants had to upload their results and technical reports. Finally, on 30 June 2025, the challenge results and rankings were published in the website. BioDCASE 2025 concluded with a workshop on the 29th of October at Universitat Pompeu Fabra, Barcelona, hosted as a satellite event to DCASE (30-31st October). The workshop featured two invited talks related to bioacoustics and spotlight presentations and posters from task participants.

We next elaborate on the three tasks carried out during the BioDCASE 2025 challenge.

### 2.3. BioDCASE 2025 tasks

#### 2.3.1. Task 1: Multi-channel audio alignment

Scientists often deploy multiple audio recorders simultaneously, for example with passive automated recording units (ARU’s) or embedded in animal-borne bio-loggers. Analysing sounds simultaneously captured by multiple recorders can provide insights into animal location and abundance (Malinka et al., 2020), caller identity (Zeh et al., 2024), as well as the dynamics of communication in groups (Gill, Goymann, Ter Maat, & Gahr, 2015). However, these devices are susceptible to desynchronisation due to non-linear clock drift (Anisimov et al., 2014). Motivated by this, Task 1 asked participants to develop a post-processing-based resynchronisation method, in order to increase usability of desynchronised collected data. To our knowledge, this task was relatively unexplored, with only one previous publication (Smeele, Tyndel, Klump, Alarcón-Nieto, & Aplin, 2024) focused on general-purpose bioacoustic resynchronisation; others, e.g. Malinka et al. (2020), employ a timing signal logged by recording devices to enable resynchronisation. The space of possible solutions was therefore relatively unexplored, with ample opportunity for participants to contribute novel methods.

Participants were asked to design a system that, when presented with pairs of temporally desynchronised recordings, could synchronise them in time (Figure 1). In the development phase, participants were provided audio pairs and a small set of ground-truth synchronisation key points–the likes of which could be produced by a manual review of the data. In the evaluation phase, participants’ systems were ranked by their ability to synchronise audio pairs drawn from the same recording session. Each dataset provided to participants consisted of a number of 115-second stereo audio recordings, where the two channels were desynchronised in time. For the development set, participants also were provided with a text file of ground-truth keypoints.

**Figure 1.**
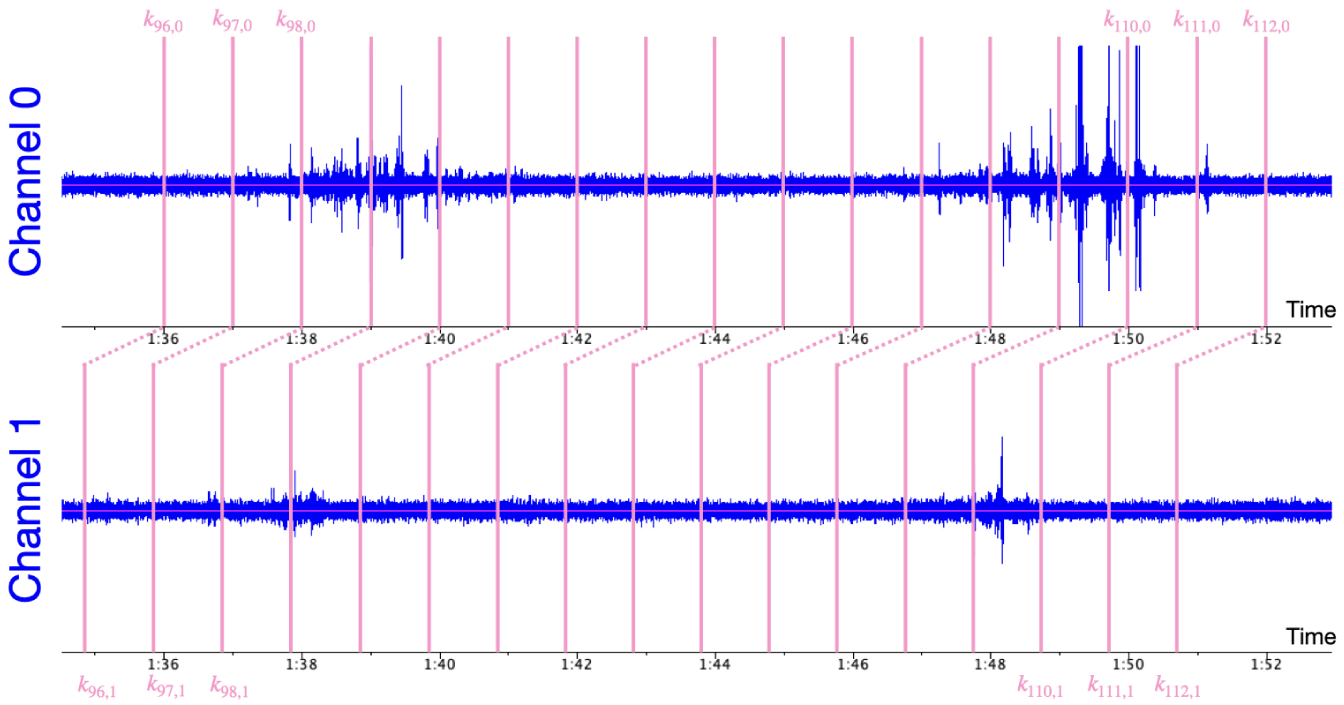
Keypoint-based alignment in task 1. The audio in the two channels of each audio file were not synchronised in time. Each audio file had a set of 115 keypoint annotations *k*_0_*, …, k*_114_. Each keypoint *k_i_*= (*k_i,_*_0_*, k_i,_*_1_) consists of a timestamp *k_i,_*_0_ for Channel 0 and a timestamp *k_i,_*_1_ for Channel 1. The timestamps in each channel correspond to the same time in the physical world, but due to clock drift they do not occur at the same time in the recordings. Systems for task 1 were provided with the *k_i,_*_0_ timestamps, and had to predict the *k_i,_*_1_ timestamps.

**Datasets:** Two multi-channel datasets were used, in order to reflect typical situations in which desynchronisation may occur. The first dataset, aru, consisted of ARU recordings made in forests in the USA and UK. The second dataset, zebra finch, consisted of on- and off-body recordings made of zebra finches (*Taeniopygia guttata*) in a laboratory setting by Gill et al. (2016). To generate both challenge datasets, we began with source recordings that were exactly synchronised in time: the microphones in the aru recordings were wired to a single recorder, and the microphones in the zebra finch recordings transmitted captured audio to a single recorder with AM radio waves. For each pair of simultaneous recordings, we introduced desynchronisation using custom signal processing transformations (we do not disclose them at this time, because the challenge may run again in the future). This approach allowed us to know the exact resynchronisation required for each pair of recordings, enabling precise measurement of the error in each participant’s proposed solution.

A key element of this task was that there were acoustic events that were present in one recording, but not the other. This occurred when the microphones were spread apart, with only one microphone was close enough to the sound source to record it. In this situation, methods that rely purely on acoustic similarity for resynchronisation, such as the one proposed by Smeele et al. (2024), may ascribe a low similarity score to the correct alignment.

**Baselines:** Three baseline systems of varying complexity were provided, which participants could reference and modify when designing their own submissions. The first of these baselines, Nosync, did not attempt to perform any synchronisation; it reflected the error present in the provided desynchronised clips. The second, Crosscorrelation Baseline, was loosely based on the approach of Smeele et al. (2024). This method found the alignment between recordings which maximised spectral cross-correlation across the two audio clips. It was a purely signal-processing-based approach and did not rely on the development set for tuning parameters.

The final, DLBaseline, used a binary classifier that is trained to determine, for a pair of 1-sec mono audio clips, whether they are aligned in time or are not. The model takes two 1-sec mono audio clips as input, and outputs either 1 (the clips are aligned in time) or 0 (the clips are not aligned in time). For each clip, audio features are extracted using a frozen pre-trained BEATs encoder. The features from each clip are averaged in time, and then the time-averaged features from the two clips are concatenated. The concatenated features are passed through a multi-layer perceptron (MLP), which is trained using binary cross-entropy loss. The classifier was trained separately on each of the two challenge datasets.

To use the DLBaseline classifier to produce the keypoint predictions for an entire audio file, candidate keypoint sets are generated under the assumption that the desynchronisation between channels consists of a constant offset plus linear time drift. Each candidate alignment is divided into 1-sec windows; the trained classifier then predicts if each of these windows is aligned in time. An alignment is scored based on the number of 1-sec windows which are predicted to be aligned. The candidate set of keypoints with the highest score is chosen as the final alignment prediction for that audio file.

#### 2.3.2. Task 2: Supervised detection of strongly-labelled Antarctic blue and fin whale calls

In underwater PAM (henceforth UPAM), the development and evaluation of generalisable machine-learning solutions, as intended in BioDCASE, have lagged slightly behind that of terrestrial PAM. Over the past decade, the taxonomic and geographic diversity represented in the recordings of the Watkins database (Sayigh et al., 2016) has strongly motivated its adoption for such research studies. Noticeably, its best-cut portions have been incorporated into the BEANS benchmark (Hagiwara et al., 2023), which in turn has been employed in multiple investigations of model generalisation (Burns et al., 2025; van Merriënboer et al., 2025). Also, the UPAM community has already implemented initiatives in the area of data challenges, though at smaller scales than neighbouring disciplines. The DCLDE workshop (https://www.dclde2024.com/previous/) has been organising a data challenge since its first edition in 2003, albeit without a systematic focus on direct competitive evaluation of multiple systems. Nonetheless, these biennial workshops have resulted in peer-reviewed publications or special journal issues (e.g. Adam et al. (2006)), and the availability and persistence of the challenge datasets can still produce a ‘long-tail’ of publications beyond that of the workshop (Hildebrand et al., 2022; Shiu et al., 2020; Sugarman, Ferguson, Alongi, Schallert, & Lyn, 2025).

We aimed to build on these efforts with goals of enhancing community engagement across machine learning, marine science, and bioacoustics communities; improving rigour in assessing performance of solutions with respect to the specific application requirements of UPAM; and providing a widely applicable, contemporary, and forward-looking benchmark for the community that is rooted in open-science foundations. We further sought to provide an accessible and well-documented entry point for both new and experienced model developers interested in UPAM. Through careful selection of training and evaluation datasets for our challenge we aimed to explore the ideas of generalisation, particularly with regard to domain shift in UPAM.

Lastly, from the complementary perspective of ocean scientists, policy makers, and managers operating within UPAM-based frameworks, our BioDCASE initiative is intended to address the need for coordinated, collaborative efforts focused on generating operationally relevant observations that are suitable for integration into global ocean observing programs, such as GOOS (Tyack et al., 2023).

**Dataset and evaluation framework:** In UPAM, generalisation is often considered in terms of well-defined factors such as sites, time periods, or recording platforms. The “hold-out dataset” approach constitutes a direct and practical implementation of such scenarios, i.e. holding entire datasets in reserve for evaluation. Thus, Task 2 asked participants to develop and test automatic detectors and classifiers on different datasets collected across multiple locations around Antarctica over several years (Miller et al., 2021).

Collected and annotated by the ICW-SORP/SOOS Acoustic Trend Working Group, these datasets comprise recordings from 11 “site–year” pairs. The annotation effort focused on calls from Antarctic blue whales (*Balaenoptera musculus*) and fin whales (*Balaenoptera physalus*), resulting in seven labels (four blue whale–specific, two fin whale–specific, and one common to both species). Owing to the characteristics of these calls, the original seven classes were grouped into three final labels for the task. The overall objective was to design model(s) able to detect sound events and to classify them into these three classes. During the development phase, the 11 site–year pairs were split into training and validation sets, both curated and made available on Zenodo^2^. Totalling 1,880 hours of audio distributed across 6,500 audio files, these datasets contained over 76,000 manually annotated sound events. Annotations were provided in CSV files (one per site-year pair) as time–frequency bounding boxes. The frequency component was included as “extra” information, as the final evaluation was based solely on one-dimensional intersection over union (IoU, i.e. the ratio of the temporal overlap between the prediction and the ground truth to the total duration defined by the earliest start time and the latest end time) along the temporal axis. During the evaluation phase, participants were provided with a test set comprising two new site–year pairs that were never made public before, without associated annotation files. It consisted of 408 audio files, totalling 400 hours of audio containing over 12,000 sound events. The developed models were then applied to these data and ranked based on their ability to detect and classify sound events using one-dimensional IoU (see subsection 3.2 for results).

The use of 1D IoU to evaluate the models was part of the task-specific considerations to be addressed. This choice was motivated by the fact that the outputs of automatic detectors may be used for downstream tasks such as density estimation, for which obtaining an accurate number of calls is crucial. Not only does IoU enable this, but it is also customisable, allowing model outputs to be penalised or rewarded by selecting which detections are taken into account for the final call count. In the context of this task and the need for accurate call counts, the custom IoU was designed to penalise models that produced multiple detections for a single ground-truth event: only the first detection was counted, while subsequent ones were discarded as false positives. Another consideration that arose during task design concerned the training/validation/test split among the 13 datasets ultimately used. On the one hand, the goal was to emphasise model generalisation ability, such that training data overly similar to validation or test data would be detrimental; on the other hand, introducing an unbridgeable domain shift would have discouraged participants and rendered the task nearly impossible. Assuming that changes in recording location and/or year would be sufficient to assess generalisation, the choice was therefore made to introduce only one novelty in the validation set (one dataset was recorded during a year unseen in the training set) and two in the test set (one site present in both training and validation data but from a year not present in any of them, and one site–year pair that was entirely unseen by both training and validation set).

Finally, alongside the model development task, a re-annotation campaign of the evaluation sets was launched. Using the APLOSE annotation platform (Dubus et al., 2025), users with different levels of expertise (expert, intermediate, or novice) were provided with spectrograms to annotate, spanning both test sets. The objective of this campaign was to study multi-annotator setups, to investigate what multi-annotator variability reveals about the data, and to compare model performance across different annotation contexts (Dubus et al., 2024; Duc et al., 2021; Leroy, Thomisch, Royer, Boebel, & Van Opzeeland, 2018).

**Baselines:** To provide both introductory material and reference benchmarks, we proposed two baseline models for our task: YOLOv11 and ResNet18. Prior approaches have been proposed in the literature, including signal-processing based methods (Miller et al., 2021; Schall & Parcerisas, 2022) and deep learning approaches (Miller, Madhusudhana, Aulich, & Kelly, 2023; Schall, Kaya, Debusschere, Devos, & Parcerisas, 2024); we deliberately selected well-known and readily accessible models in order to lower the barrier to entry and facilitate reproducibility.

YOLOv11 is a computer vision model with support for object detection, segmentation, classification, and more. It is one of the latest iterations in the YOLO (You Only Look Once) series, improving the models performance in several ways such as enhanced feature extraction with the introduction of the C3k2 (Cross Stage Partial with kernel size 2) block, SPPF (Spatial Pyramid Pooling - Fast), and C2PSA (Convolutional block with Parallel Spatial Attention) components (Khanam & Hussain, 2024). For the YOLOv11 baseline, recordings were chunked into windows larger than the longest expected sound event of interest and converted to spectrograms. The present events in the windows were re-annotated as 2D bounding-boxes, in a similar way to the common YOLO computer vision tasks. The model was trained directly using the ultralytics package with default values, providing room for improvement for the participants. Data-augmentation approaches which are not relevant for acoustics data were deactivated.

The ResNet-18 model is the smallest and most lightweight variant of the original residual network (ResNet) architecture (He, Zhang, Ren, & Sun, 2016). Consisting of 18 learnable layers (mainly convolutional and one fully connected layer), it contains significantly fewer parameters than deeper ResNet variants, making it relatively easy to train and deploy, hence making it a perfect starting point. Unlike YOLO, which is an object detection model, ResNet is primarily designed for image classification, producing a vector of class probabilities indicating the likelihood of each category being present in the image. This output format constrains the inputs to also be vector of probabilities. To do so, audio files were chunked into smaller pieces that got re-annotated, using binary vectors for each of the three labels mentioned above (zero-vectors were considered “noisy” chunks) and were then fed to the model. Its implementation, along with its training and evaluation was made using the torchvision library.

Source code and documentation can be found on Github^3^. During the model development phase, we additionally conducted a comprehensive baseline evaluation on both the training^4^ and validation^5^ datasets. On the validation set, YOLOv11 attained an F1 score of 0.43, exceeding the F1 score of 0.32 achieved by the ResNet-based model. The performance gap can be explained by differences in supervision and spatial resolution. Indeed, ResNet eventually compresses inputs into a single global representation via global average pooling, which removes fine-grained temporal and frequency information. Trained on coarse audio chunks, it only predicts event presence within a segment. In contrast, YOLO is trained with precise annotation boundaries and operates at a finer resolution, allowing accurate localisation of sound events in time–frequency space.

#### 2.3.3. Task 3: Bioacoustics for tiny hardware

For the sake of ecology and conservation research, there is a need for long-term deployments of acoustic sensors often in remote locations, and almost always off the electrical grid. Hence a challenge for computational bioacoustics: how to design sensors that are able to operate reliably over long deployment periods despite infrequent energy supply and maintenance? The current generation of sensors are certainly lighter and more energy-efficient than before. Yet, despite them being presented as “autonomous recording units” (ARUs), several obstacles come in the way of truly autonomous passive acoustic monitoring:

**Downtime.** Storing hundreds of hours of audio data onto an SD card incurs a significant expense of energy. Current-generation hardware lowers this expense by resorting to an intermittent acquisition schedule, which may come at the detriment of detectability for rare acoustic events (Teixeira, Maron, & van Rensburg, 2019).

**Maintenance costs.** When travelling to remote sites, batteries are heavy, and subject to travel restrictions, which adds to the practical difficulty of fieldwork (Wood et al., 2023). Further, travelling costs associated with revisiting sites for maintenance make a large part of monitoring costs even when travelling locally (Markova-Nenova, Engler, Cord, & Wätzold, 2023).

**Invasiveness.** Frequent round trips between the base station and deployment sites generate noise and other anthropogenic disturbances (Vallee, 2018).

**Surveillance creep.** Because some terrestrial datasets contain voices from people who are unaware of being recorded (Cretois, Rosten, & Sethi, 2022), the same data which are collected for research may be used for surveillance (Parker & Dockray, 2023).

**Environmental and social harms.** The use of batteries is linked to labour rights violations during mineral extraction; carbon emissions during production and transport; toxicity to ecosystems including humans; and the risk of improper handling of e-waste (Melchor-Martínez et al., 2021; Sharma & Manthiram, 2020).

Task 3 was designed to reflect realistic constraints encountered when deploying bioacoustic detection: namely, memory footprint, computational complexity, and latency. The challenge consisted in developing a real-time, on-device vocalisation detection system for the Yellowhammer (*Emberiza citrinella*) running on the ESP32-S3-Korvo-2 microcontroller. The motivation for this task was to record all interactions of Yellowhammers at the location over the breeding season from March to August. Similar tasks are common in conservation practice or ecological research since conservation managers or researchers often need to know the presence and activity of target species in the area.

**Datasets:** The development set consisted of 2-second audio clips of Yellowhammer songs, amounting to 2 hours and 37 minutes in total. The dataset originates from a sound transmission experiment which consisted of playing a track combining 209 high-quality songs from 10 individuals (21 song types, approximately 10 repetitions each) in forest and grassland environments and recording the playback at seven distances ranging from 6.5 to 200 m using AudioMoth recorders. The recordings were then manually annotated and the clips resampled to 16 kHz. The dataset was split by individual into training (6), validation (2), and evaluation (2) sets, negative samples were randomly assigned.

**Baseline:** To lower the entry barrier and ensure fair benchmarking, a comprehensive end-to-end baseline framework was provided (Carmantini, Förstner, Isik, & Kahl, 2025), covering data preprocessing, feature extraction implemented consistently on both host and device, model training, automated deployment and benchmarking on the target hardware. Customisation was permitted at two key stages: during feature extraction, where participants could tune parameters to optimise their approach, and during model training, where they could define custom architectures and, if needed, implement a custom training loop. The baseline framework served both as a starting point for participants unfamiliar with embedded ML development and as a reference implementation demonstrating the expected integration between model development on the host and deployment and execution on the embedded device.

During the evaluation phase, participants’ models were assessed by the organisers on the hidden evaluation set using the baseline framework’s benchmarking tools. Models were ranked based on metrics capturing both classification performance and resource efficiency.

## 3. Results

The first edition of the challenge attracted a diverse set of participants across its three tasks. For Task 1, a total of five participant submissions were received, in addition to the three baseline systems provided by the task organisers. Task 2 received twelve participant submissions, along with the official baseline system. Finally, Task 3 received seven participant submissions. The quantitative metrics of the three tasks are summarised in Tables 1, 2, and 3. Each table report the performance of the teams according to the official evaluation metrics defined for each task, which are further discussed in Subsections 3.1, 3.2 and 3.3.

**Table 1.** Task 1: model summary performance per submission. Sorted by rank score.

| Name | Rank Score | Test Set |  | Validation Set |  |  |
| --- | --- | --- | --- | --- | --- | --- |
|  | MSE avg<br>(Test) | MSE<br>(aru) | MSE<br>(Zebra Finch) | MSE<br>(avg) | MSE<br>(aru) | MSE<br>(Zebra Finch) |
| BEATsCA | 0.30 | 0.14 | 0.45 | 0.31 | 0.10 | 0.52 |
| DLBaseline | 0.58 | 0.62 | 0.55 | 0.89 | 0.52 | 1.26 |
| CP-MedianLag | 1.29 | 0.36 | 2.22 | 3.00 | 0.26 | 5.74 |
| CP-MedianLag_Regress | 1.33 | 0.23 | 2.43 | 2.96 | 0.66 | 5.26 |
| Nosync Baseline | 1.35 | 0.85 | 1.84 | 1.15 | 0.98 | 1.31 |
| Landmarks | 1.60 | 1.60 | 2.04 | 0.84 | 0.41 | 1.27 |
| Crosscorrelation<br>Baseline | 3.32 | 1.10 | 5.55 | 8.45 | 6.86 | 10.03 |
| CP-GlobalLag | 3.93 | 0.62 | 7.23 | 5.30 | 0.39 | 10.20 |

**Table 2.** Task 2: model summary performance per submission. Sorted by rank score.

| Name | Rank Score | Dataset metrics |  | Overall metrics |  |  |
| --- | --- | --- | --- | --- | --- | --- |
| Metric (%) |  | DDU2021<br>F-Score | Kerguelen2020<br>F-Score | Overall<br>F-Score | Overall<br>Recall | Overall<br>Precision |
| Swin transformer fusion with softmax | 50.00 | 46.40 | 50.40 | 50.00 | 47.70 | 52.50 |
| Swin transformer fusion with softmax | 43.80 | 38.80 | 53.40 | 43.80 | 33.80 | 62.40 |
| Baseline YOLOv11 | 39.20 | 39.50 | 42.70 | 39.20 | 33.10 | 48.00 |
| Whale-VAD | 35.70 | 33.90 | 36.30 | 35.70 | 38.20 | 33.60 |
| Whale-VAD | 34.90 | 34.20 | 32.10 | 34.90 | 41.40 | 30.20 |
| Whale-VAD | 33.60 | 33.90 | 31.90 | 33.60 | 45.80 | 26.60 |
| Tiny yolov4 coco | 27.50 | 27.50 | 28.90 | 27.50 | 17.50 | 63.70 |
| Voxaboxen dv | 19.20 | 20.50 | 11.20 | 19.20 | 18.10 | 20.60 |
| Voxaboxen dv | 17.80 | 18.50 | 14.10 | 17.80 | 13.90 | 24.90 |
| Voxaboxen dv | 15.90 | 16.40 | 13.20 | 15.90 | 10.80 | 29.90 |
| Enhanced Spectrogram-YOLO | 8.60 | 7.20 | 16.00 | 8.60 | 4.60 | 70.80 |
| Edge optimised CNN | 8.50 | 7.60 | 14.00 | 8.50 | 4.80 | 35.50 |
| Edge optimised CNN | 4.40 | 3.50 | 10.60 | 4.40 | 2.40 | 31.90 |

**Table 3.** Task 3: model summary performance per submission.

| Name | Avg. Precision (%) | Model Size (B) | RAM Usage (B) | Preprocessing Time (ms) | Model Time (ms) |
| --- | --- | --- | --- | --- | --- |
| Londono_enriched | 98 | 16600 | 26844 | 176.11 | 24.42 |
| Londono_non_enriched | 94 | 16600 | 26844 | 176.22 | 24.42 |
| Christian_Walter | 74 | 7020 | 6744 | 1.62 | 16.45 |
| Oguamanam_team | 97 | 4192 | 18354 | 34.28 | 7.39 |
| Oguamanam_team_flux | 97 | 1848 | 780 | 16.54 | 0.18 |
| Toby_Martin | 99 | 7776 | 20924 | 73.42 | 15.16 |
| Naveen_Dhar | 74 | 12920 | 30328 | 64.84 | 112.78 |

In terms of the workshop, 113 attendees (including volunteers) registered prior to the event. Among them, 94 provided their nationalities, representing 24 different countries as shown in Figure 2. The figure illustrates the broad international reach of the event, with a notable participation in Spain due to physical closeness to the venue. It should be emphasised that not all attendees were participants in the BioDCASE challenge; a number of them were DCASE conference attendees who joined the workshop as researchers with a general interest in the topic. The workshop program consisted of: (1) two invited talks of 45 minutes of duration; (2) a set of three presentations by task organisers, who introduced the objectives, datasets and evaluation metrics of the challenge tasks; (3) nine participants contributions—some of which were not related to the BioDCASE tasks but addressed broader bioacoustics topics—, consisting of brief spotlight presentation and a dedicated poster session; and (4) an open discussion and panel session focused on future directions in bioacoustics research and the long-term development of the BioDCASE initiative.

**Figure 2.**
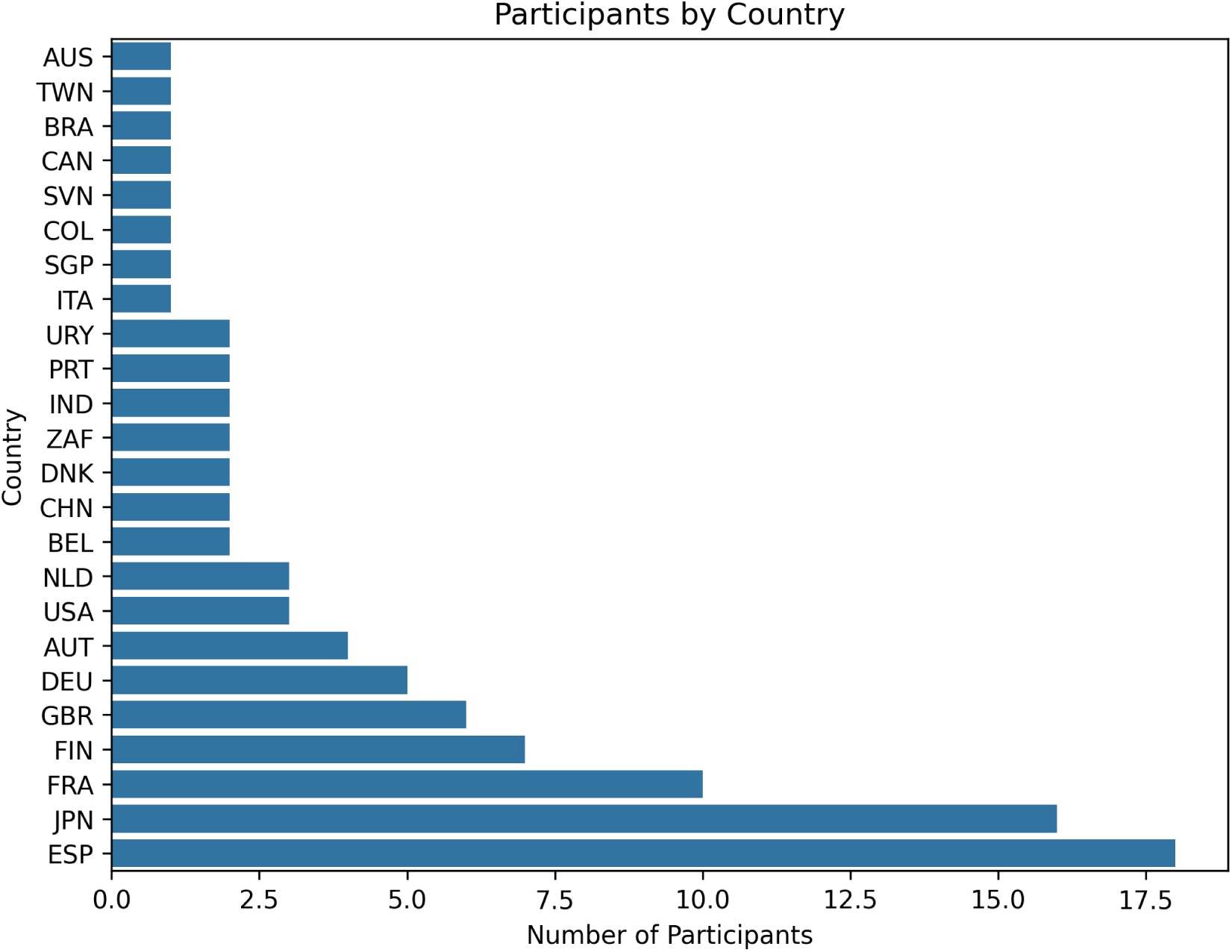
Geographical origin of the BioDCASE 2025 workshop attendees. Countries are represented with their ISO 3166-1 alpha-3 code.

### 3.1. Task 1: Multi-channel audio alignment – outcomes

Task 1 received five submissions from three teams. Nihal, Yen, Ashizawa, and Nakadai (2025) proposed the top performing system BEATsCA, which was a modification of the DLBaseline. It introduced several improvements, including 1) a head with more layers and cross-attention across the per-channel embeddings; 2) amplitude-scaling and additive noise data augmentations and 3) a custom scoring function that incorporates model confidence.

Bhattacharjee (2025) proposed the second submitted system CP-MedianLag and its variants CP-MedianLag Regress and CP-GlobalLag. Two of these out-performed the DLBaseline on the aru dataset. This system introduced contrastive pretraining on mixtures of audio clips drawn from FSD50k Fonseca, Favory, Pons, Font, and Serra (2021), and was applied to the challenge test datasets without any fine-tuning. Local offsets between the windowed single-channel recordings were estimated based on similarity of audio embeddings, and then a global offset between the two audio channels was set to be the median of these local offsets. The variants explored alternate methods of estimating alignment, including within-clip drift.

The third submitted system, Landmarks, identifies peaks in the mel spectrograms and matches nearby peaks based on a variety of criteria ((Harju & Mesaros, 2025)). A degree-3 polynomial was fit to the matched peaks, to generate the final alignment.

Overall, the results of this challenge demonstrate this the multi-channel alignment task is challenging, with high inter-dataset variability in task difficulty. Signal processing methods based on landmarks and spectral cross-correlation did not improve on the Nosync baseline. The unsupervised method CP-MedianLag showed promise on the aru dataset, but struggled to generalise to the zebrafinch dataset. Possibly, this is explained by the relative similarity of the challenge datasets to that submission’s pretraining data. The supervised approach introduced with DLBaseline and improved by BEATsCA shows promise as a general method of multi-channel alignment. Future work could improve on this method, or explore alternative supervised and unsupervised alignment strategies.

### 3.2. Task 2: Detection of Antarctic blue and fin whale calls – outcomes

For task 2 of the bioDCASE challenge 2025 on the detection of baleen whale calls, all participants used systems that had fixed-length time windows as input, transformed the waveform to a time-frequency representation and used a neural network to extract features from the time-frequency input. The neural networks were CNN or transformer based and optimised a training loss for either multi-label classification or a box detector with YOLO. For most participating systems post-processing at inference was important, including thresholding for multi-label classification systems, non-maximum suppression, or temporal merging. The first ranking team established an end-to-end learning system: using a multi-scale learnable frontend for audio classification, i.e. creating a time-frequency representation with learnable filters from the data at 3 scales, using a Swin transformer on each of these scales and fusing the outputs to get the embedding representation for the audio before a classifier layer.

#### 3.2.1. Evaluation procedure

The macro-F1 score was used to rank the model submissions according to the ratio of recall and precision. According to a recent benchmark in baleen whale acoustics, however, also false positive rates should be evaluated in order to test model applicability to real-world long-term datasets (Schall et al., 2024). The two highest ranking model submissions result in false positive rates between 0.02 and 1.79% with an average of 0.54%, which is already very close to or even below the recommendation of false positive rates not exceeding 1%. While this is a very promising outcome, all model submissions show relatively poor abilities in finding true positives, with overall recall averages of below 50%. Potential explanations for the low recall could be class imbalance, low Signal-to-Noise ratios (SNRs) of logged vocalisations in the test dataset, or possibly that models do not make use of acoustic context.

The evaluation of submissions and the official final ranking of submitted model results was based on the annotations of two single experts who each reviewed one part of the provided test dataset from two locations, one already included in the training dataset and one new location (previously unseen for all submitted models). Overall, submitted models showed good generalisation as performance on the dataset from the new location did not generally drop in comparison to performance on the dataset from the known location.

#### 3.2.2. Using inter-annotator agreement for more robust evaluation

One potential strategy for improving the performance of automated systems is to use multiple annotators, and to filter annotations for inter-annotator-agreement, reducing subjective variation. We conducted an initial exploration of this using the preliminary results from the multi-annotator campaign, which was run in parallel to the BioDCASE 2025 challenge, simply by exploring the effect of applying this technique to the test set. The performance of the submitted models increased by on average 2% (range: −6% to +11%) when performance is measured against two expert annotators that have to agree with each other, in comparison to the performance calculated from the annotations of a single expert.

### 3.3. Task 3: Bioacoustics for tiny hardware – outcomes

The Bioacoustics for Tiny Hardware task received seven submissions from 5 different teams, which were evaluated using four metrics: (1) total execution time in milliseconds, including both preprocessing and model inference; (2) model file size (in bytes); (3) peak memory usage (in bytes); and (4) average precision. These metrics were selected to capture the trade-offs between detection performance and computational or time constraints: the first three primarily assess each model’s suitability for on-device deployment and efficiency under limited resources, while the last metric evaluates detection accuracy.

Most submissions improved significantly over the baseline (×2 up to ×29 speed up and 48% to 98% less RAM usage with under 6% loss in performance in most cases). Within these improvements, the submissions exhibited a wide range of performance–efficiency trade-offs (Table 3, Figure 3). Greater variation was observed in execution time, while average precision remained relatively consistent across solutions, aside from a few cases with unusually low precision, some of which likely resulted from difficulties in running and evaluating them using the developed framework. This variability reflects the diversity of solutions for this task:

**Figure 3.**
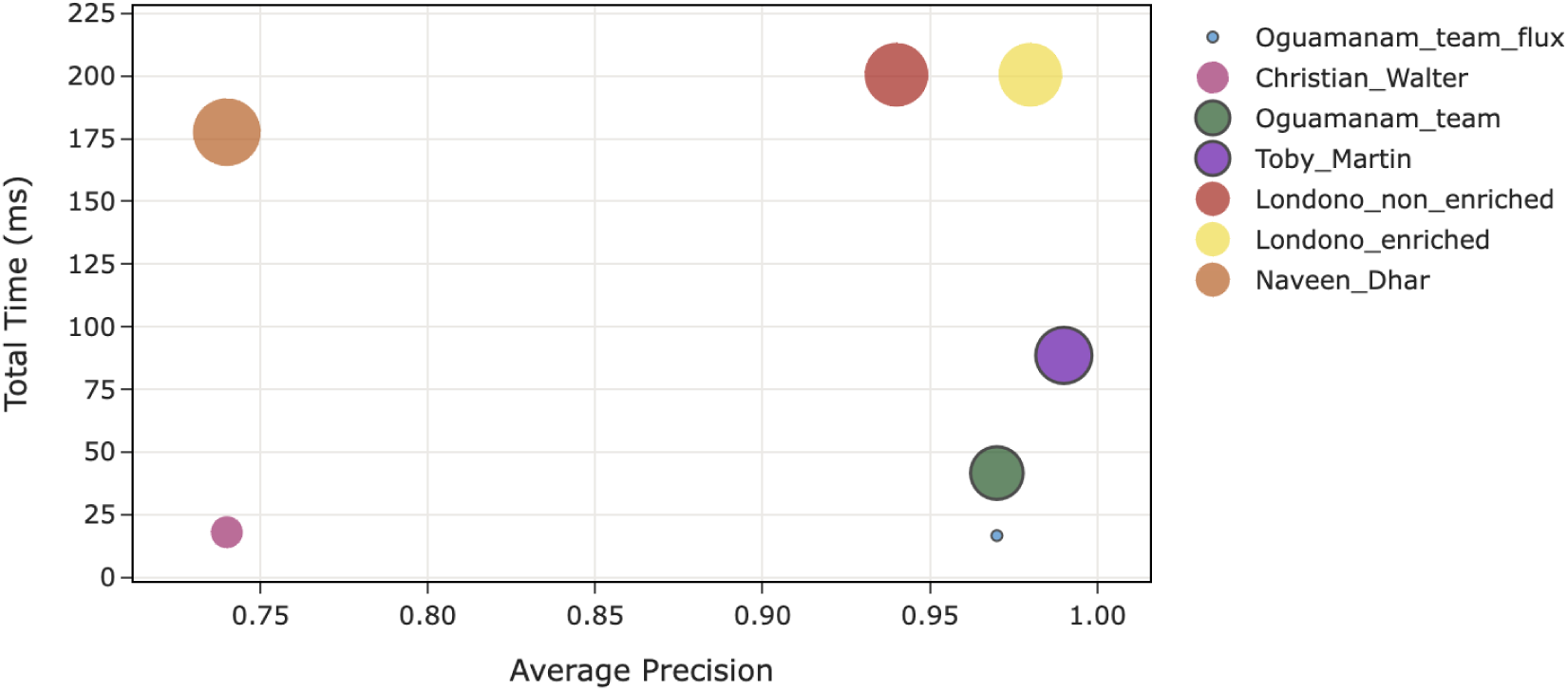
Results of Task 3 showing average precision versus total execution time, with marker size indicating RAM usage. The three solutions highlighted in the bottom-right region appear to achieve the most favourable trade-off between performance and efficiency.

**Data augmentation.** some participants employed data augmentation to enhance model robustness across different signal-to-noise ratio conditions. In particular, pitch shifting, noise addition, and SpecMixup were used, with some submissions applying these transformations dynamically according to SNR estimates.

**Feature engineering and preprocessing.** Log-mel spectrograms were the predominant input representation. (One team investigated gammatone frequency cepstral coefficients and spectral flux features.) Parameter settings, including window size, number of filters, frequency range and sample rate varied across submissions and were directly reflected in the processing time. Nevertheless, in most submissions, preprocessing still dominated the total execution time, often exceeding 80%.

**Model design and optimisation.** a broad range of model architectures and optimisation approaches were investigated. Some participants focused on architectural and training optimisations, followed by post-training quantisation, while others used techniques such as knowledge distillation, magnitude-based pruning, and quantisation-aware training. Architecturally, teams explored everything from very shallow CNNs to highly simplified MobileNet-inspired networks and convolutional–recurrent hybrids. Notably, one of the top-performing solutions used support vector machines (SVMs).

The first edition of this task provided an open-source baseline that lowered the barrier to entry and enabled straightforward end-to-end development of Tiny bioacoustic AI on the ESP32-S3-Korvo-2 microcontroller. However, difficulties in running some submissions on-device highlighted limitations in the current submission process, indicating that the submission system—and potentially the evaluation metrics—may need to be rethought.

Outcomes suggest that real-time, on-device single-species detection is feasible even under severe resource constraints and across a broad range of feasible design strategies. Recording only relevant sound data lies at the core of efficient sensor design and it has multiple benefits. For many tasks, on-device detection represents way towards greater sensor autonomy. It allows increasing uptime with limited resources, especially for rarely vocalising species. Further, it saves maintenance, post-processing and storage costs and avoids surveillance creep. However, the current task and dataset may not have been sufficiently challenging, as most submissions achieved very good performance. Future editions should consider incorporating more complex datasets and tasks to further explore innovative design strategies and allow for a deeper investigation of the performance–efficiency trade-off.

## 4. Discussion

The BioDCASE 2025 initiative was overall successful. The three tasks set clear goals with associated public data sets, there was good community uptake and involvement. In each of the tasks clear comparative evaluation was performed with multiple candidate solutions.

Key lessons were learnt from this initiative which will inform future iterations. Notably, the task difficulty level is an important aspect, dependent on the state of the art and the dataset characteristics—both of which are factors usually not fully understood before the event. The difficulty should be high enough to incentivise advances in technique, but low enough that fruitful participation is possible. The results indicate that Task 1 may have been more difficult than expected—it proved difficult to outperform the baselines—while Task 3 easier than expected, with very high precision for almost all submitted systems. Following similar experiences in the DCASE initiative (Mesaros et al., 2025), we consider it useful to repeat and refine challenge tasks over multiple editions.

Some of the issues raised in our evaluation are common issues in applied machine learning or acoustics, which will stubbornly persist due to the nature of the domain. Detection tasks such as Task 2 are often characterised by a very high class imbalance, and this remains a persistent issue for data-driven methods. Conditions of low signal-to-noise ratio (SNR) also remain challenging, seen in Task 2 but will persist in almost any bioacoustic monitoring in the wild. Another challenge is the issue of generalisation. The task evaluation should be designed such that evaluation scores provide meaningful estimates of an algorithm’s performance in wider conditions subject to domain-shift. This remains an open-challenge relevant to the evaluation of any data-driven method in computational bioacoustic.

To structure the discussion we next consider separately the opportunities and the limitations of this data challenge format, in the context of computational bioacoustics and its applications.

### 4.1. Opportunities

First, as with DCASE and many other data challenges, we observe that BioDCASE has served as an accessible entry-point to computational bioacoustics. This is important for newcomers and students, of course, but it gains extra importance due to the cross-disciplinary nature of the topic: even experienced researchers may benefit from this accessibility.

Cross-disciplinary collaboration is a key benefit we see in BioDCASE. Participation across domains such as computer science and ecology bridges expertise gaps and fosters understanding of each domain’s issues. In particular, the data challenge format helps by bringing confusions to the surface: in this format, open questions must be crystallised into explicit tasks with clear success criteria. In our forum terrestrial and marine tasks are addressed within the same space, bridging these communities. There remains much more scope for this bridging: even within the limits of the three tasks we conducted, we find that terrestrial bioacoustics could benefit from audio synchronisation work in marine applications, while marine bioacoustics stands to benefit from the generalisable deep learning work already explored in terrestrial bioacoustics.

BioDCASE tasks are open for participation from both academia and industry, and in 2025 we had many private companies taking part. A data challenge can act as a public forum for matching competencies and finding appropriate companies and contacts. Even though the data challenge is driven by a spirit of open science, this does not exclude private enterprise.

One subtle aspect is the level of computational difficulty in the tasks. Our goal is to make computational advances in method, not just to solve empirical issues. This is a higher level of computational ambition than some Kaggle challenges whose audience may be more general data scientists or students. BioDCASE task design can generate interesting tasks for computer scientists, and engage computer science at the state-of-the-art or higher academic level.

To date we have run only one iteration of BioDCASE, although with some inheritance from the regular DCASE challenge. In this context, the annual incremental improvement of task design is valuable. It is not necessarily expected that any given issue will be solved in one year, and aspects of task design—such as the difficulty level or the provided datasets—can be refined progressively. For this, the public community orientation helps to share lessons learnt. Tasks could be open to submission of additional datasets or accompanied with calls for specific data scrutinising the computational solutions. Multi-year tasks also allow to address tasks from different perspectives developing the tasks over the time towards generalised solutions or targeting different domain specific challenges.

Further, BioDCASE has a role as an umbrella platform for multiple different tasks. This helps to provide the critical mass of community size, when a task on its own might not. This also occurs initiatives such as CLEF, a long-running evaluation series within which LifeCLEF emerged (Goëau et al., 2016, 2014).The question of whether such initiatives should merge together, or retain a separate focus, is a contextual issue of community organisation.

Community evaluation initiatives can also lead to the creation of tools and resources that have long-term impact. Public datasets are a clear example. We also wish to highlight evaluation toolkits, such as the ‘sed eval’ from DCASE (Mesaros, Heittola, & Virtanen, 2016), or the ‘BioDCASE-tiny’ toolkit developed in our Task 3.

### 4.2. Limitations and challenges

A public data challenge is just one type of initiative. It is worth identifying the limitations in such an approach. We begin with general limitations that have been identified, before considering limitations specific to BioDCASE and its particular cross-disciplinary context.

Firstly, the focus on clear quantitative evaluation is beneficial, but has side effects. The ranking of submissions can encourage a competitive aspect and focus attention only on the highest-rated entries. It can also encourage work that is narrowly focussed on optimising the chosen performance measure rather than on original principled research. To offset this risk, we advocate our practice of making a jury award that is more holistic.

There remains a strong reliance on datasets to drive challenge tasks. In some cases, though not always, it is desired to have new unreleased data each year, which creates a continuous demand for data. Some bioacoustic needs which are not typically reflected in large annotated datasets—such as vocal interactions or spatial details—must be addressed by other means. Further, although some datasets are large, they are still limited in balance, representation and in inter-annotator consistency. Within Task 2 we explored this latter issue and determined its impact could be significant. However, we acknowledge that multi-annotation of any large bioacoustic dataset is often hard to justify the in person-time required.

Code availability of proposed solutions is not enforced, which hinders the adoption of solutions by others or iteratively improving upon it. It does however enable participation by others including private companies. To mitigate, task organisers provide open code for baselines, and can also incentivise—e.g. openness being a requirement for the official awards.

Forming new cross-disciplinary teams and collaborations is indeed a goal of BioDCASE, but also a challenging goal. Considering especially that BioDCASE addresses an audience with diverse skills brought from ecology, computer science, acoustics and so forth, it might be valuable to offer some skill-matching team formation function. This is not currently part of the programme, meaning that teams typically are pre-existing collaborators, or must find connections on their own. The success of networks across all tasks, at generalisation across datasets is often constrained by a lack of deep learning technical ability (ecological teams) vs a lack of deep understanding of the biological variance within datasets, ocean soundscapes and drivers of ambient diversity (computer scientist entries). To drive forward innovation in this field at a pace equal to that of hardware challenges, there should be opportunities for cross-field collaboration to inject expertise from all aspects of the pipeline. This applies to task organising teams as much as to participant teams.

We have already referred to the required balance of designing tasks that are general, while also having concrete practical relevance for bioacoustic practitioners. We consider the niche of BioDCASE to be aiming towards the general, which leaves room for other initiatives to develop work focussed on more pragmatic use-cases.

### 4.3. Developing good tasks in computational bioacoustics

There are many possible issues to solve in computational bioacoustics. It may not always be clear which of those are amenable to the format of public data challenge. It is therefore appropriate to consider the attributes of a successful task, and how they might relate to open issues that have been identified in the literature. Many of these derive directly from our design principles; others arise from experience.

**Broad Relevance.** A task should represent a broad category, for which multiple groups might be able to use the solution. This is key to findings having relevance to the broader computational bioacoustics community.

**Accessible.** Tasks must be designed to be sufficiently accessible by participants. This means balancing the scope and complexity of the challenge with other considerations. A specific example that has emerged in DCASE is the exclusion of ‘sub-tasks’ which create additional complexity and risk subdividing attention.

**Measurable.** A task should have a measure of success: usually, one or more quantitative evaluation metrics. This does not rule out tasks that are qualitative or interactive in nature, but care is needed to ensure independent teams can work towards that measure of success. In our Task 3, teams typically developed their solutions on virtual (emulated) hardware. The emulator helps to make the task accessible. However, final evaluation was on-device, which was subtly different.

**Generalisable.** Task results should generalise beyond the scope of the challenge. It is important that evaluation data captures the complexity of real-world deployment conditions (e.g. domain-shift, SNR, imbalance etc). Practically, this may mean including both in- and out-of-domain evaluation datasets across different recording conditions, locations, species etc.

**Data Availability.** A task should be represented by relevant dataset(s) that can be shared. This also extends to other resources such as software code and evaluation measures. Most often, the limiting factor in framing a challenge task is the range of data that are available or even possible.

- Rare species and rare sound events: There is widespread need for systems that perform well on both very common and very rare sounds, yet rarely-occurring sounds (e.g. from rare or cryptic species) are inherently sparse in datasets. This makes it very difficult to curate appropriate datasets, and to evaluate reliably. One potential avenue for this, testable in a data/algorithm-centric setting, is the use of soundscape generators to create synthetic training data with tuneable characteristics (Guei, Christin, Lecomte, & Hervet, 2023; Weldy et al., 2025).
- Conversely, some very broad categories can be difficult to build a representative dataset for—such as “anthropogenic noise”, which encompasses speech, footsteps, laughter, traffic, gunshots, shipping and much more.
- Interactive uses, such as data interfaces or active learning. Interactive tools are useful because they make the most of computation as well as human skills. Active learning alternates between human annotation and machine prediction, and this semi-automation can yield good results faster than the standard methods of supervised learning (Kath, Serafini, Campos, Gouvêa, & Sonntag, 2024; Kholghi, Phillips, Towsey, Sitbon, & Roe, 2018). But evaluating such systems is non-trivial: should evaluation include a human in the loop? Or can a fixed dataset be used to emulate a human decisionmaker?
- There are many other topics of high interest which are difficult simply because well-annotated datasets are hard to obtain. Examples include counting the number of individuals in a dense sound scene (Navine, Camp, Weldy, Denton, & Hart, 2024), or spatial aspects such as the distance from the calling animal to the recording device. These metadata could be provided in specialised circumstances, but for many datasets captured in the wild, are unknown.

In future, more bioacoustic tasks may be addressed through the use of so-called ‘foundation models’—a generic term for broadly-trained deep learning models reusable for many tasks (Schwinger et al., 2025). Such models are no silver bullet: their robust generalisation or reusability remains largely an empirical matter. We believe the principles of task design that we have discussed hold valid even as foundation models become established. Public evaluation remains valuable, and it is also likely that BioDCASE solutions (i.e. the specific engineering required to address any given task) remain valuable.

### 4.4. Conclusions

We advocate for data challenges as a method to bring bioacoustics communities together and make verifiable improvements on computational tasks that can benefit many of us. Data challenges such as BioDCASE sit alongside other traditional evaluation methods and educational initiatives, and their level of difficulty can be calibrated to the needs of the community. The most significant investments required for a BioDCASE challenge are data curation, and community facilitation.

Beyond the challenge, we have seen that datasets often become benchmarks for related tasks (Hagiwara et al., 2023; Kao & Liu, 2025; Wu, Xu, Wei, & Long, 2023). These dispersed benefits further enhance the community value of conducting such work in the paradigm of open science.

## Acknowledgments

This work was supported by the European Union’s Marie Skłodowska-Curie Action under grant agreement No 101116715 “Bioacoustic AI”; and Biodiversa+ under grant “TABMON”. PL has been supported by SustainScapes – Center for Sustainable Landscapes under Global Change (grant NNF20OC0059595). The authors report there are no competing interests to declare.

We thank other members of the task organisation who are not otherwise listed as authors (see https://biodcase.github.io/challenge2025/), as well as data annotators, and challenge participants.

## CRediT STATEMENT

Conceptualisation: DS, IN, EV-V, BM, VL. Methodology: DS, EV-V, IN, ES. Software: BH, CP. Validation: BH, ES, CP. Investigation: DS, IN, EV-V, BM, BH, ES, IMor, CP. Data Curation: LJL, GD, BH, PL, CP, IMor, YB. Writing - Original Draft: DS, EV-V, BM, IN, BH, EW, DC, PYR, ES, IMor, PL, VL, YB. Writing - Review & Editing: DS, EV-V, IN, EW, IM, PL, DC. Supervision: DS, PL.

## Footnotes

1 https://zenodo.org/

2 https://zenodo.org/records/15092732

3 https://github.com/marinebioCASE/task2_2025/tree/main/baselines

4 https://drive.google.com/file/d/17hX3S9woRBmTw2tUHsS__vgn42dB8OMf/view

5 Table 2 in https://biodcase.github.io/challenge2025/task2

## Notes

https://biodcase.github.io/challenge2025/

https://github.com/marinebioCASE/

https://zenodo.org/records/15092732

### Competing Interest Statement

The authors have declared no competing interest.

